# Cotranslational folding of human growth hormone *in vitro* and in *Escherichia coli*

**DOI:** 10.1101/2022.08.08.503169

**Authors:** Daphne Mermans, Felix Nicolaus, Aysel Baygin, Gunnar von Heijne

## Abstract

Human growth hormone (hGH) is a four-helix bundle protein of considerable pharmacological interest. Recombinant hGH is produced in bacteria, yet little is known about its folding during expression in *E. coli*. We have studied the cotranslational folding of hGH using Force Profile Analysis (FPA), both during *in vitro* translation in the absence and presence of the chaperone trigger factor (TF), and when expressed in *E. coli*. We find that the main folding transition starts before hGH is completely released from the ribosome, and that it can interact with TF and possibly other chaperones.

## Introduction

Human growth hormone (hGH) is used world-wide to treat a number of conditions, including dwarfism, bone fractures, and skin burns [1, 2]. Because un-glycosylated hGH is biologically active and the protein is not subject to any other post-translational modification critical for its activity, it is commonly produced intracellularly in bacteria and purified for pharmaceutical use [3–5].

*In vitro*, purified hGH typically reaches the fully folded state on a timescale of ∼3 sec via a very rapid collapse phase, and the folding rate is independent of the formation of disulfide bonds [6]. This suggests that hGH may start to fold cotranslationally when produced in bacteria. Further, it is not known whether cytoplasmic chaperones such as trigger factor (TF) may impact the folding of hGH in bacteria.

In order to address these questions, we have probed the cotranslational folding of hGH, both in *E. coli* and in the *E. coli*-derived PURExpress™ *in vitro* transcription-translation system, using Force Profile Analysis (FPA). FPA takes advantage of so-called translational arrest peptides (APs) – short stretches of polypeptide that bind with high affinity in the upper reaches of the ribosome exit tunnel and thereby arrest translation at a specific codon in the mRNA [7]. The translational arrest can be overcome if a strong enough pulling force is exerted on the AP, essentially pulling it out of its binding site in the exit tunnel [8–11]. APs can be employed as sensitive “molecular force sensors” to report on various cotranslational events such as protein folding [12–16], protein translocation [17], and the membrane protein integration [9,18,19].

We find that hGH undergoes cotranslational folding the *in vitro* system shortly before the entire protein has emerged from the ribosome exit tunnel. The presence of TF delays the onset of folding by ∼4 residues, suggesting that hGH interacts with TF during translation. A similar delay is seen *in vivo*, consistent with chaperone interactions during the folding of hGH.

## Materials and methods

### Key Resources Table

| *Key Resources Table* |  |  |  |  |
| --- | --- | --- | --- | --- |
| *Reagent type (species) or resource* | *Designation* | *Source or reference* | *Identifiers* | *Additional information* |
| Strain, strain background (*Escherichia coli*) | MC1061 | J Biol Chem. 261:13844-9. PMID: 3531212 | NA | Electrocompetent cells |
| Other | Protein G-Agarose | Roche | 11243233001 | Resin used for immunoprecipitation |
| Antibody | Anti-HA.11 Epitope Tag Antibody (mouse monoclonal) IgG | BioLegend | Cat# 901533 | Used for immunoprecipitation (1 µl of 1 mg/mL, diluted 1:820) |
| Recombinant DNA reagent | pET-19b (plasmid) | Novagen | Cat # 69677 | Expression plasmid |
| Recombinant DNA reagent | pING1 (plasmid) | Gene 34:137-45. PMID: 4007491. | NA | Expression plasmid |
| Commercial assay, kit | GeneJET Plasmid miniprep kit | Thermo Fisher Scientific<br>RRID:SCR_008452 | Cat no. 0502 | Used to purify plasmids |
| Commercial assay, kit | GeneJET PCR Purification Kit | Thermo Fisher Scientific | Cat no. K0701 | Used to purify linear fragments for *in vitro* expression |
| Commercial assay, kit | PURExpress | New England Biolabs | Cat no. E6800L | Used for *in vitro* expression |
| Chemical compound, drug | <sup>35</sup>S Methionine | Perkin-Elmer | Cat no. NEG009T001MC | <sup>35</sup>S Methionine is incorporated into the protein during *in vitro* and *in vivo* translation and aids detection by phosphorimaging. |
| Software, algorithm | EasyQuant | Developed in-house<br>Nat Struct Mol Biol. 19:1018-22.<br>PMID: 23001004; |  | Used to quantify relative fraction full-length of translated protein from SDS-PAGE |

### Enzymes and chemicals

All enzymes used in this study were purchased from Thermo Scientific and New England Biolabs. Oligonucleotides were from Eurofins Genomics. DNA isolation/purification kits and precast polyacrylamide gels were from Thermo Scientific. L-[^35^S]-methionine was obtained from PerkinElmer. Mouse monoclonal antibody against the HA antigen was purchased from BioLegend. Protein-G agarose beads were manufactured by Roche. All other reagents were from Sigma-Aldrich.

### Cloning and Mutagenesis

Various hGH constructs followed by a variable LepB-derived linker sequence (between 4 and 34 residues), the 9-residue long HA tag, the 17-residue long *E. coli* SecM AP, and a 23-residue long C-terminal tail were engineered in pING1 (used for *in vivo* expression) and pET-19b (used for *in vitro* expression) plasmids. The hGH constructs were cloned into the respective plasmids using Gibson assembly^®^ [20].

For the *in vitro* assays, ordered gene fragments were used to introduce the C-terminal GS-repeats in construct *N* = 182. Point mutations and C-terminal truncations were done by performing site-specific DNA mutagenesis. All cloning and mutagenesis products were confirmed by DNA sequencing. hGH sequences used in this study are summarized in Supplementary File 1.

### In vivo pulse-labeling analysis

Competent *E. coli* MC1061 cells [21] were transformed with a pING1 plasmid, and grown overnight at 37°C in M9 minimal medium supplemented with 19 amino acids (1μg/ml, no Met), 100 μg/ml thiamine, 0.4% (w/v) fructose, 100 mg/ml ampicillin, 2 mM MgSO_4_, and 0.1mM CaCl_2_. Cells were diluted into fresh M9 medium to an OD_600_ of 0.1 and grown until an OD_600_ of 0.3-0.5. Expression was induced with 0.2% (w/v) arabinose for 5 minutes at 37°C followed by [^35^S]-Met pulse-labeling (2 minutes). The reaction was stopped using 20% ice-cold trichloroacetic acid (TCA). Samples were put on ice for 30 min and immunoprecipitation using an antibody directed against the HA tag was carried out as previously described [19]. Samples were precipitated, washed and immediately solubilized in SDS sample buffer before incubation with RNase and analysis by SDS-PAGE.

Radiolabeled proteins were detected by exposing dried gels to phosphorimaging plates, which were scanned in a Fuji FLA-3000 scanner. Band intensity profiles were obtained using the FIJI (ImageJ) software and quantified with our in-house software EasyQuant. *A_c_* and/or *FL_c_* controls were included in the SDS-PAGE analysis for constructs where the identities of the *A* and *FL* bands were not immediately obvious on the gel. Data was collected from at least three independent biological replicates, and averages and standard errors of the mean (SEM) were calculated. Statistical significance was calculated in Microsoft Excel using a two-sided t-test.

### In vitro transcription/translation

*In vitro* coupled transcription-translation was performed in PURExpress™ (New England Biolabs, Ipswich, MA, USA), according to the manufacturer’s instructions, using PCR products as templates for the generation of truncated nascent chains. Briefly, 1 μl (50 ng) of PCR template and 10 μCi (1 μl) [^35^S]-Methionine were added to a final volume of 10 μl reaction components. The chaperone trigger factor (TF) was added to the reaction mixtures to a final concentration of 4 μM. Transcription-translation was carried out for 15 min at 700 rpm in a bench-top tube shaker at 37°C. Reactions were stopped by adding equal volumes of ice-cold 10% TCA, incubated on ice for 30 min., and then centrifuged at 14,000 rpm (20,800 x g) at 4°C for 10 min. in an Eppendorf centrifuge. Pellets were resuspended in 30 μl of SDS Sample Buffer and treated with RNase A (400 μg ml^−1^) for 15 min. at 37 °C. Proteins were separated by SDS-PAGE, visualized on a Fuji FLA-3000 phosphorimager, and quantified using the FIJI (ImageJ) software. Analysis of quantified bands was performed using EasyQuant (in-house developed quantification software). Values of *f*_FL_ were calculated as *f*_FL_ = *I*_FL_/(*I*_FL_ + *I*_A_), where *I*_FL_ is the intensity of the band corresponding to the full-length protein, and *I*_A_ is the intensity of the band corresponding to the arrested form of the protein. Experiments were repeated at least three times, and SEMs were calculated.

## Results

### Force Profile Analysis

Translational arrest peptides (APs) are short stretches of polypeptide that interact with the ribosome exit tunnel in such a way that translation is stalled when the ribosome reaches the last codon in the AP [22]. The stall can be overcome by external forces pulling on the nascent chain [23], and the stalling efficiency of a given AP is reduced in proportion to the magnitude of the external pulling force [8, 9]. APs can therefore be used as force sensors to follow a range of cotranslational processes that generate force on the nascent chain such as membrane protein biogenesis [9, 24], protein translocation [17], and protein folding [12,13,15,25–27].

A schematic representation of how cotranslational protein folding generates force on the AP is shown in Fig. 1a. For short constructs for which there is not enough room in the ribosome exit tunnel for the protein to fold at the point when the ribosome reaches the end of the AP, or for long constructs where the protein has already folded when the ribosome reaches the end of the AP, little force is generated and translation is efficiently stalled. However, for constructs of intermediate length where there is just enough space in the tunnel for the protein to fold if the tether is stretched out from its equilibrium length, some of the free energy gained upon folding will be stored as elastic energy (increased tension) in the nascent chain, reducing stalling. By measuring the stalling efficiency for a series of constructs of increasing length, a force profile (FP) can be generated that shows how the folding force varies with the location of the protein in the exit tunnel [15], and hence when during translation the protein starts to fold. The approach has been validated in a number of ways. It has been shown that the main peak in a FP correlates with the acquisition of thermolysin-resistance of the folding protein in an on-ribosome pulse-proteolysis assay [12], and with the appearance of folded protein in the exit tunnel as visualized by cryo-EM [13,15,26]. Further, the amplitude of the peak correlates with the thermodynamic stability of the folded protein [12]. Finally, FPs can be quantitatively reproduced by molecular dynamics simulations of cotranslational protein folding [15, 26], and are affected by changes in the size and shape of the exit tunnel in ways expected for cotranslational folding [27, 28].

**Figure 1.**
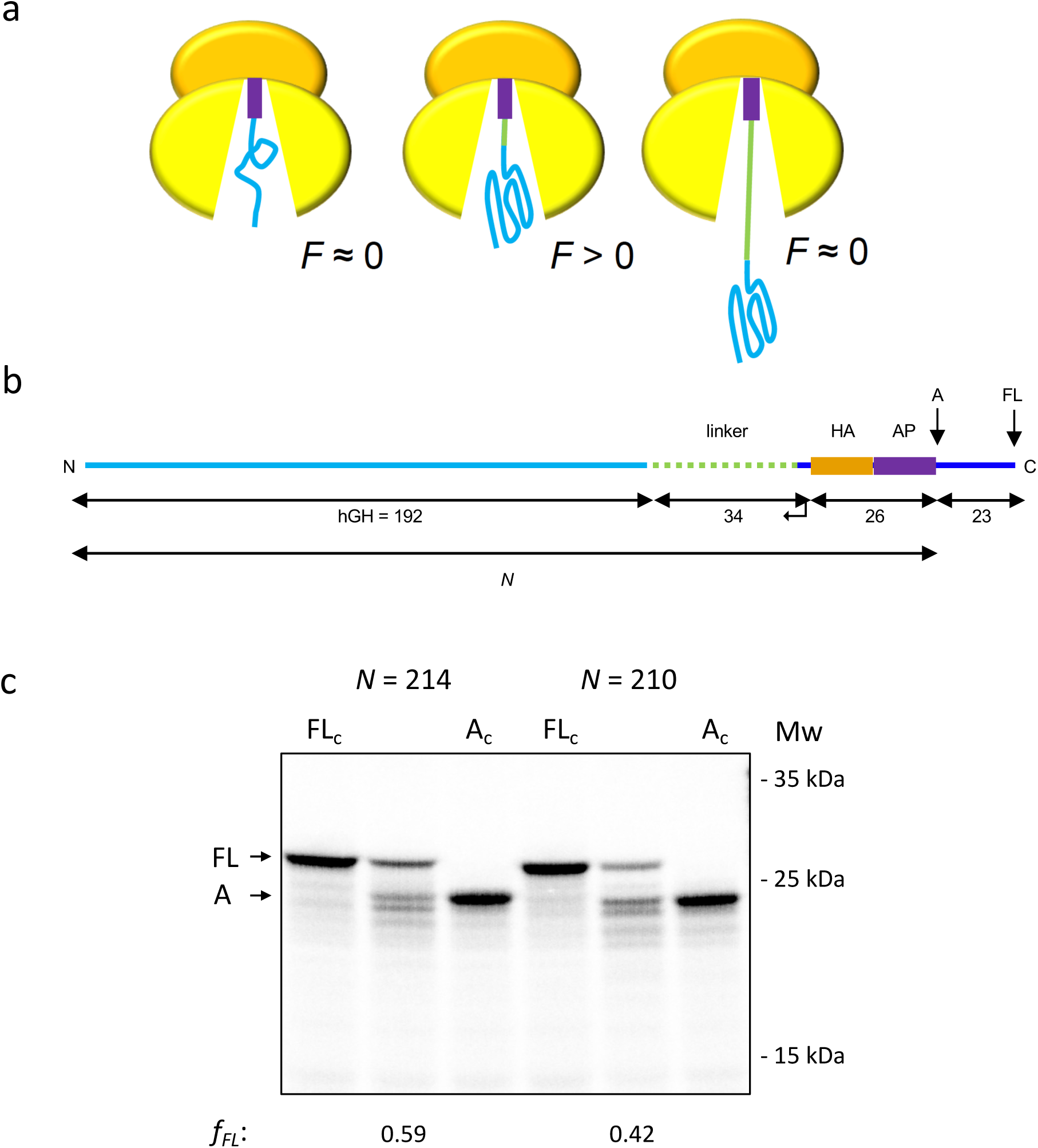
The force profile assay. (a) Schematic scenario for constructs generating (*F* > 0) or not generating (*F* ≈ 0) pulling force depending on the location of hGH domain (in blue) relative to the AP (in purple). (b) Basic construct. hGH, variable linker, HA-tag, SecM(*Ec*) AP, and C-terminal tail are indicated, together with the arrested (*A*) and full-length (*FL*) products. *N* denotes the number of residues between the N-terminal end of hGH and the last residue (Pro) in the AP. (c) SDS-PAGE gels for constructs *N* = 214 and *N* = 210. Full-length (FL) and arrested (A) products are indicated. FL_c_ is a full-length control where the AP has been inactivated by mutation of the C-terminal Pro residue to Ala; A_c_ is an arrest control when the codon for the last Pro in the AP has been mutated to a stop codon. The band immediately below the A product likely results from ribosome stacking behind the stalled ribosome.

The basic construct used to obtain hGH FPs is shown in Fig. 1b (note that we count the initiator Met as residue number one, while this residue is usually not counted in biochemical studies of purified hGH). The C-terminal end of hGH is connected, via a variable-length linker and a HA-tag (for immunoprecipitation), to the *E. coli* SecM AP [7], which in turn is followed by a 23-residue C-terminal tail (included to allow separation of arrested and full-length forms of the protein by SDS-PAGE). Constructs are translated for 15 min. in the PURExpress™ coupled *in vitro* transcription-translation system [29] in the presence of [^35^S]-Met, the radiolabeled protein products are analyzed by SDS-PAGE, and bands are quantitated on a phosphoimager, Fig. 1c. For constructs where little pulling force is exerted on the AP, stalling is efficient and the arrested (A) form of the protein dominates. In contrast, for constructs experiencing a high pulling force there is little stalling and the full-length (FL) form dominates. We use the fraction full-length protein, *f_FL_* = *I_FL_*/(*I_FL_*+*I_A_*) (where *I_i_* is the intensity of band *i* = A, FL), as a measure of the force exerted on the AP [9]. The DNA and protein sequences of all constructs analyzed are given in Supplementary File 1.

### hGH starts to fold when its C-terminus is ∼20 residues away from the PTC

In a first set of experiments, we measured a FP for hGH folding *in vitro*, using PURExpress™, a coupled transcription-translation system manufactured from purified *E. coli* components. As seen in Fig. 2a, there is a major peak in the FP that reaches half-maximal amplitude at *N_onset_* ≈ 212 residues, *i.e*., when the C-terminal F^192^ in hGH is ∼20 residues away from the peptidyl transferase center (PTC) in the ribosome. To ascertain whether this peak reflects folding of hGH, we mutated two aromatic residues, W^87^ and F^167^, in the hydrophobic core of the protein to Ala, Fig 2b, in an attempt to destabilize the folded state. Indeed, both mutations significantly reduced *f_FL_* at *N* = 218 residues, Fig, 2c, while a control mutation (F^98^A) in a location outside the central core had no effect on *f_FL_*. Likewise, mutation of either of the two C-terminally located Cys residues to Ala had no effect on *f_FL_*. We thus conclude that this peak reflects the folding of hGH as it emerges from the exit tunnel.

**Figure 2.**
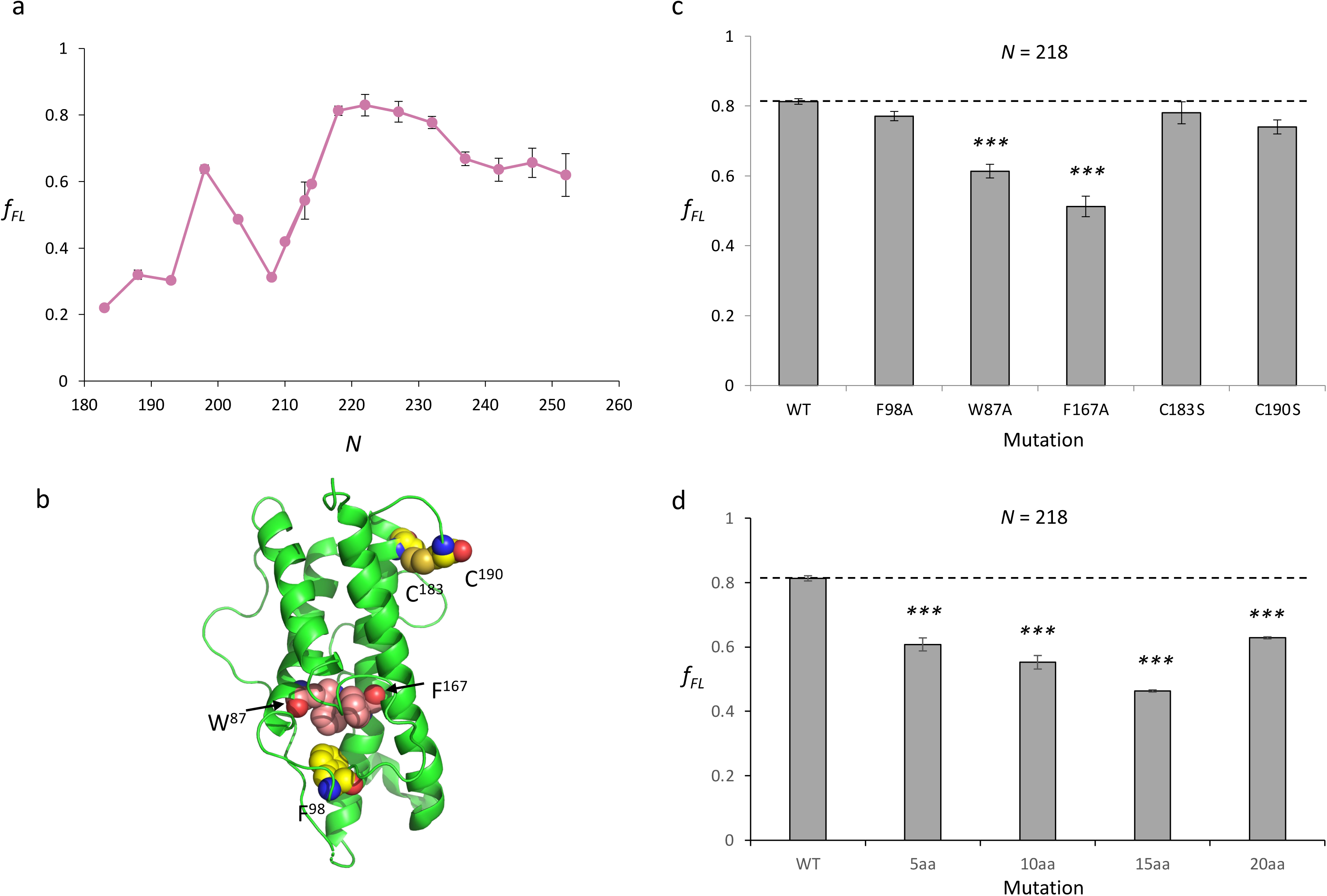
hHG FPs recorded in the PURExpress™ in vitro translation system. (a) hGH FP. (b) hGH with mutated residues indicated in spacefill (PDB 1HGU). (c) *f_FL_* values at *N* = 218 for the indicated hGH mutants. (d) *f_FL_* values at *N* = 218 for constructs in which 5, 10, 15 and 20 C-terminal hGH residues have been replaced by GS-repeats. Error bars show SEM values (*n* = 3). In panels *c* and *d*, two-sided t-tests were carried out to assess the significance of the observed differences in *f_FL_* values between the *N* = 218 construct (“WT”) and mutated constructs (*p* ≤0 .01: **, *p* ≤ 0.001: ***).

We further replaced 5-20 C-terminal residues in hGH with GS-repeats in construct *N* = 218. This led to a progressive reduction in *f_FL_* values, Fig. 2d. The C-terminal end of hGH thus contributes to the stability of the folded state, although at least a part of the last α-helix can be substituted with GS-repeats without completely destabilizing the protein. Interestingly, when the 20 C-terminal residues in hGH were replaced by GS-repeats, *f_FL_* increased compared to the 15-residue replacement; in this case, the GS-repeat region plus the HA-tag and the AP comprise 46 residues, *i.e*., the GS-repeat region completely fills the lower part of the exit tunnel which may minimize the interactions between the nascent chain and the tunnel wall and thereby weaken the translational arrest [12, 30]. Further work will be required to fully understand how flexible, ‘non-interacting’ polypeptides such as long GS-repeats impact translation when located in the exit tunnel.

A peak of lower amplitude is seen at *N* = 198 residues, possibly representing the formation of a cotranslational folding intermediate lacking the last ∼20 residues of hGH. In the *N* = 198 construct, the C-terminal residue D^172^ in the truncated hGH is tethered 26 residues away from the PTC, *i.e*., in a location in the ribosome exit tunnel where small protein domains of up to ∼25 residues can fold [12]. In order to probe the structure of the putative folding intermediate we mutated individual residues in the N-terminal part of the truncated helix H4 and residues surrounding H4 in the folded full-length hGH, Fig. 3a. As seen in Fig. 3b, the only mutation that significantly reduced *f_FL_* is L^164^P in H4. Notably, the F^167^A mutation that significantly reduced *f_FL_* for in the full-length *N* = 218 construct (Fig. 2c) had no effect at *N* = 198, showing that the hydrophobic core in folded full-length hGH is not formed in the *N* = 198 construct. We can also rule out that the long segment 129-158 that precedes the C-terminal α-helix contributes to the peak at *N* = 198 residues, as replacement of residues 129-143 or 144-158 with equally long GS-repeat segments had no effect on *f_FL_*, Fig. 3b. Replacing the highly charge stretch R^168^KDMD^172^ at the C-terminal end of the truncated H4 helix with GSGSG did give rise to a significant reduction in *f_FL_*, however.

**Figure 3.**
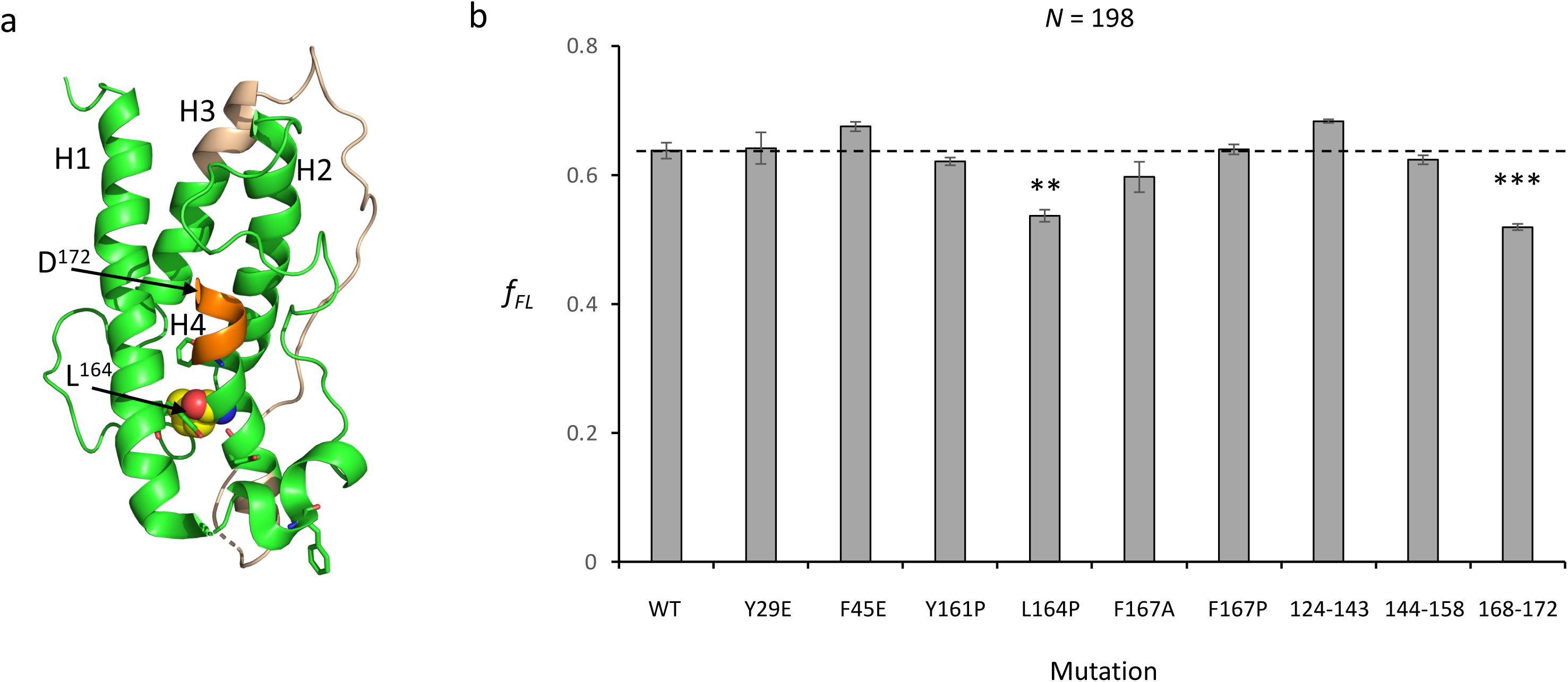
Mutational analysis of the putative folding intermediate at N = 198 residues. (a) Mutations in the *N* = 198 construct mapped onto the structure of hGH. The *N* = 198 construct is truncated at D^172^ in the middle of helix H4 which is connected to the PTC by a 26-residue linker. L^164^ is indicated in spacefill and other mutated residues in stick representation. Residues 168-172 are indicated in orange, and residues 124-158 in wheat. (b) *f_FL_* values for the *N* = 198 construct (“WT”) and for the indicated mutations. Mutations denoted “*xxx*-*yyy*” have the indicated segments replaced by GS-repeats. Error bars show SEM values (*n* = 3). Two-sided t-tests were carried out to assess the significance of the observed differences in *f_FL_* values between the *N* = 198 construct and mutated constructs (*p* ≤0 .01: **, *p* ≤ 0.001: ***).

**Figure 4.**
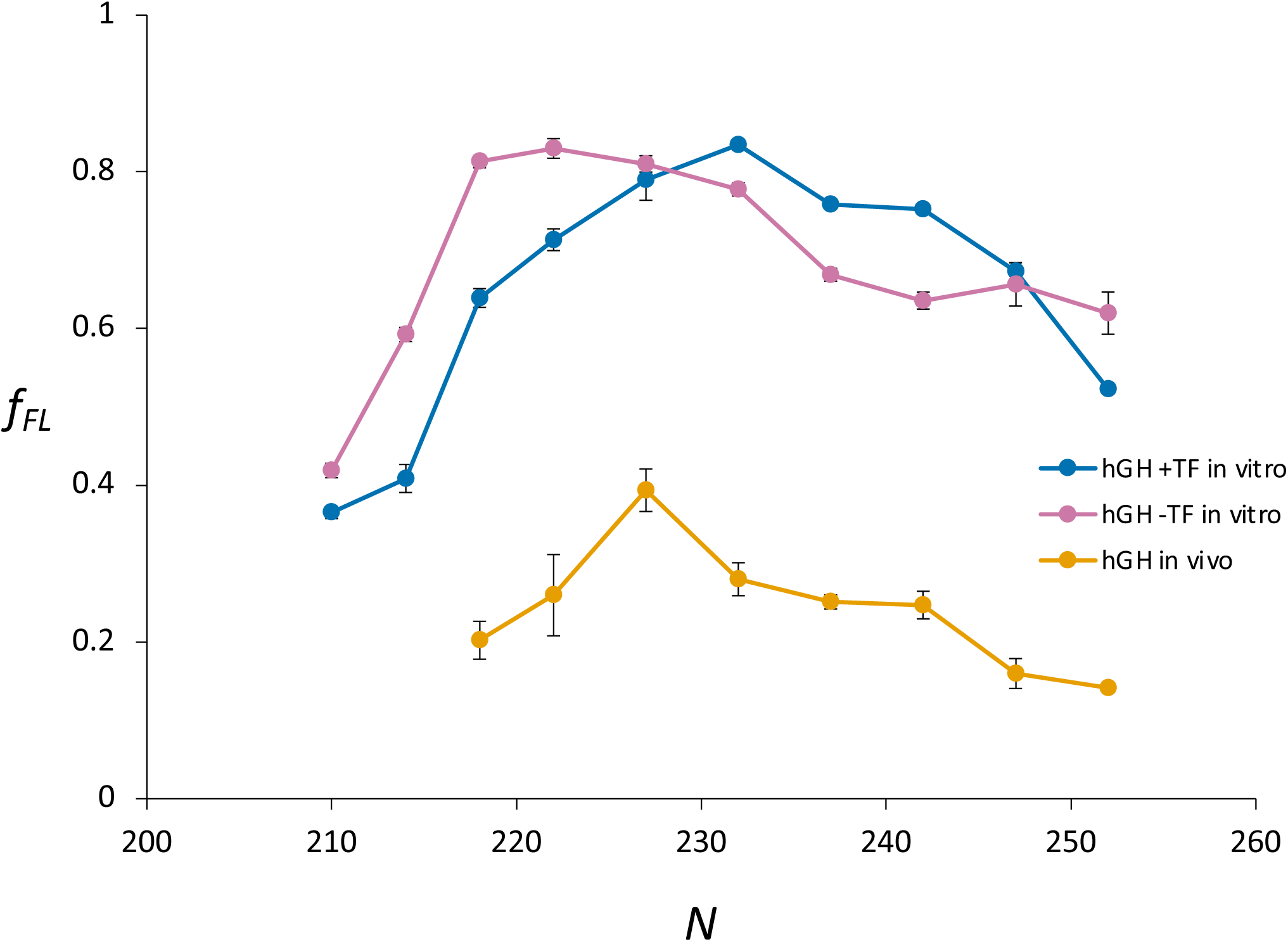
hHG FPs recorded in vivo and in the PURExpress™ system with added TF. *In vivo* hGH FP (orange), *in vitro* hGH FP (magenta; c.f., Fig 2a), and *in vitro* hGH FP recorded in the presence of 4 μM TF (blue), Error bars show SEM values (*n* = 3).

It thus appears that the peak at *N* = 198 at least in part reflects the folding of the first half of helix H4 within the exit tunnel but that the hydrophobic core in hGH (involving residues W^87^ and F^167^, Fig. 2) forms only at *N* = 218 when the C-terminal end of helix H4 emerges from the exit tunnel.

### Nascent hGH interacts with TF during cotranslational folding

To probe the role (if any) of the TF chaperone during hGH folding, we recorded an *in vitro* FP in the presence of 4 μM TF, Fig. 3 (blue curve). TF right-shifts the folding peak by ∼4 residues, demonstrating that nascent hGH interacts with TF as it emerges from the exit tunnel. This is consistent with our earlier observation of a similar TF-induced shift in the FP obtained for the cotranslational folding of DHFR, a 159-residue protein that folds outside the exit tunnel, and the absence of a shift for the 29-residue zinc-finger ADR1a that folds within the exit tunnel and hence is not available for binding to TF [25].

Finally, we recorded an *in vivo* hGH FP by radiolabeling with [^35^S]-Met for 2 min. in live *E. coli*, followed by immunoprecipitation and SDS-PAGE, Fig 3 (orange curve). The FP is again right-shifted, suggesting that, once emerged from the exit tunnel, hGH can undergo cotranslational folding and interact with cytoplasmic chaperones also *in vivo*.

## Discussion

hGH is an important pharmaceutical product that is produced in large volumes in bacteria such as *E. coli*. Yet, not much is known about the *in vivo* folding of recombinant hGH. Using FPA, we now find that hGH undergoes its main folding event shortly before its C-terminal end emerges from the ribosome exit tunnel, and that it can interact with the cytoplasmic chaperone TF during translation. Our system is artificial in the sense that we have appended a linker to the C-terminal end of hGH, meaning that the natural protein (lacking the linker) will be released from the ribosome before the main folding event takes place. Nevertheless, our data show that hGH can start to fold while still in or immediately adjacent to the ribosome exit tunnel, most likely with the participation of TF.

## Acknowledgments

Purified trigger factor was a gift from Dr. Günter Kramer, Heidelberg University, and the hGH plasmid a gift from Dr. Colin Robinson, University of Kent. This work was supported by grants from the Knut and Alice Wallenberg Foundation (2017.0323), the Novo Nordisk Fund (NNF18OC0032828), and the Swedish Research Council (621-2014-3713) to GvH, and by a Marie Curie Initial Training Network Grant (Horizon 2020, ProteinFactory 642863) to FN.

## Data availability

Gel data available on request from the authors.

---

**Supplementary File 1:**
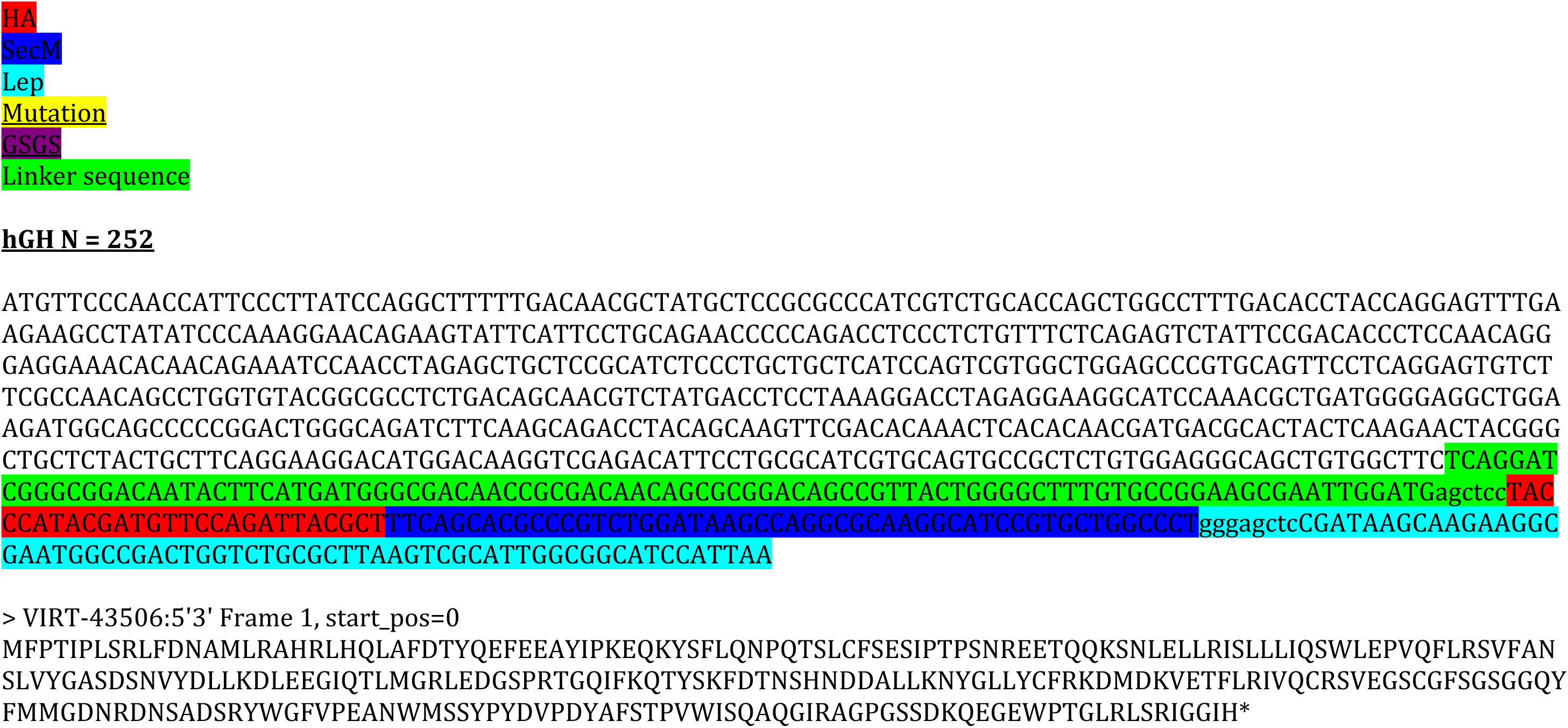
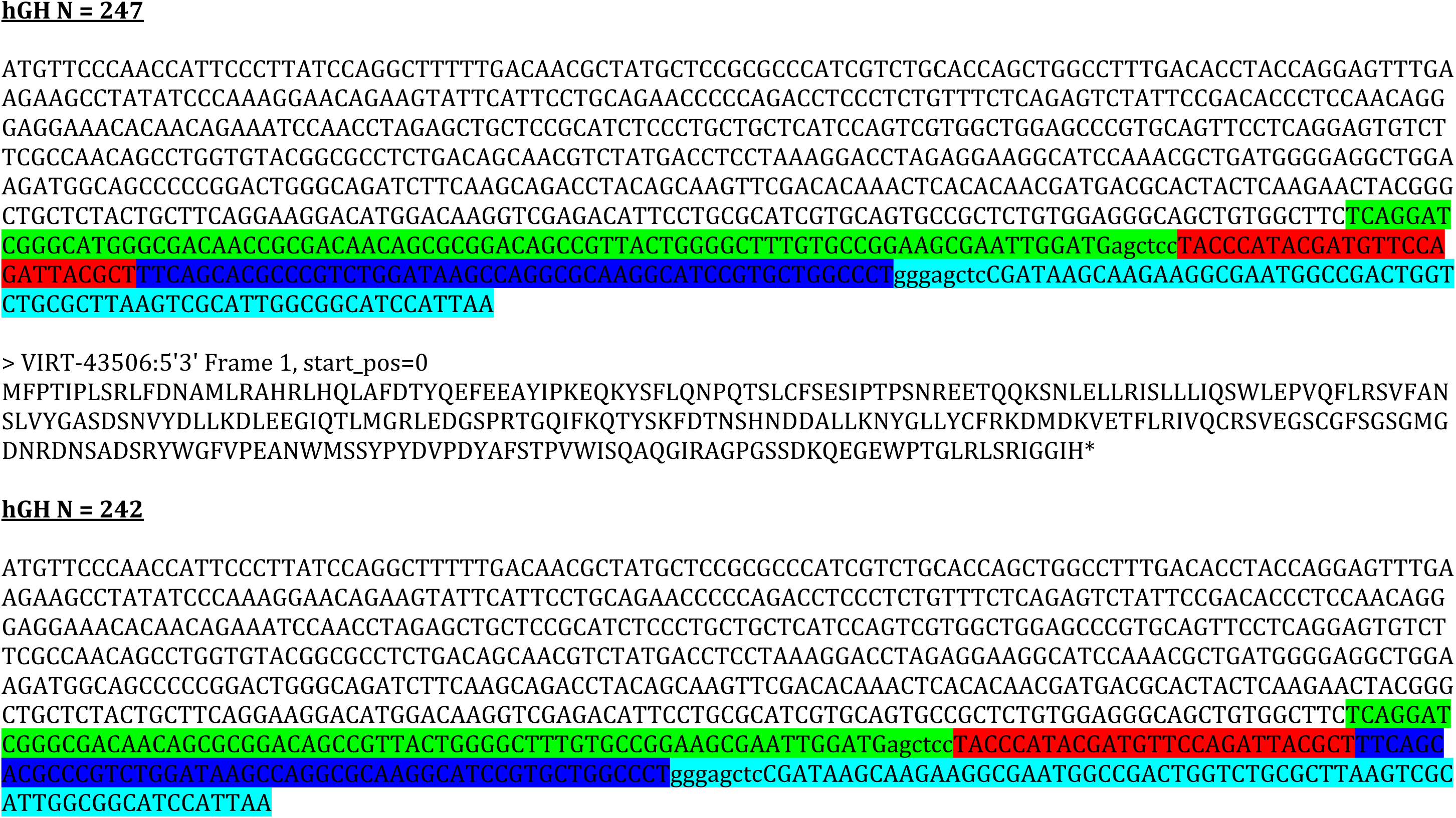
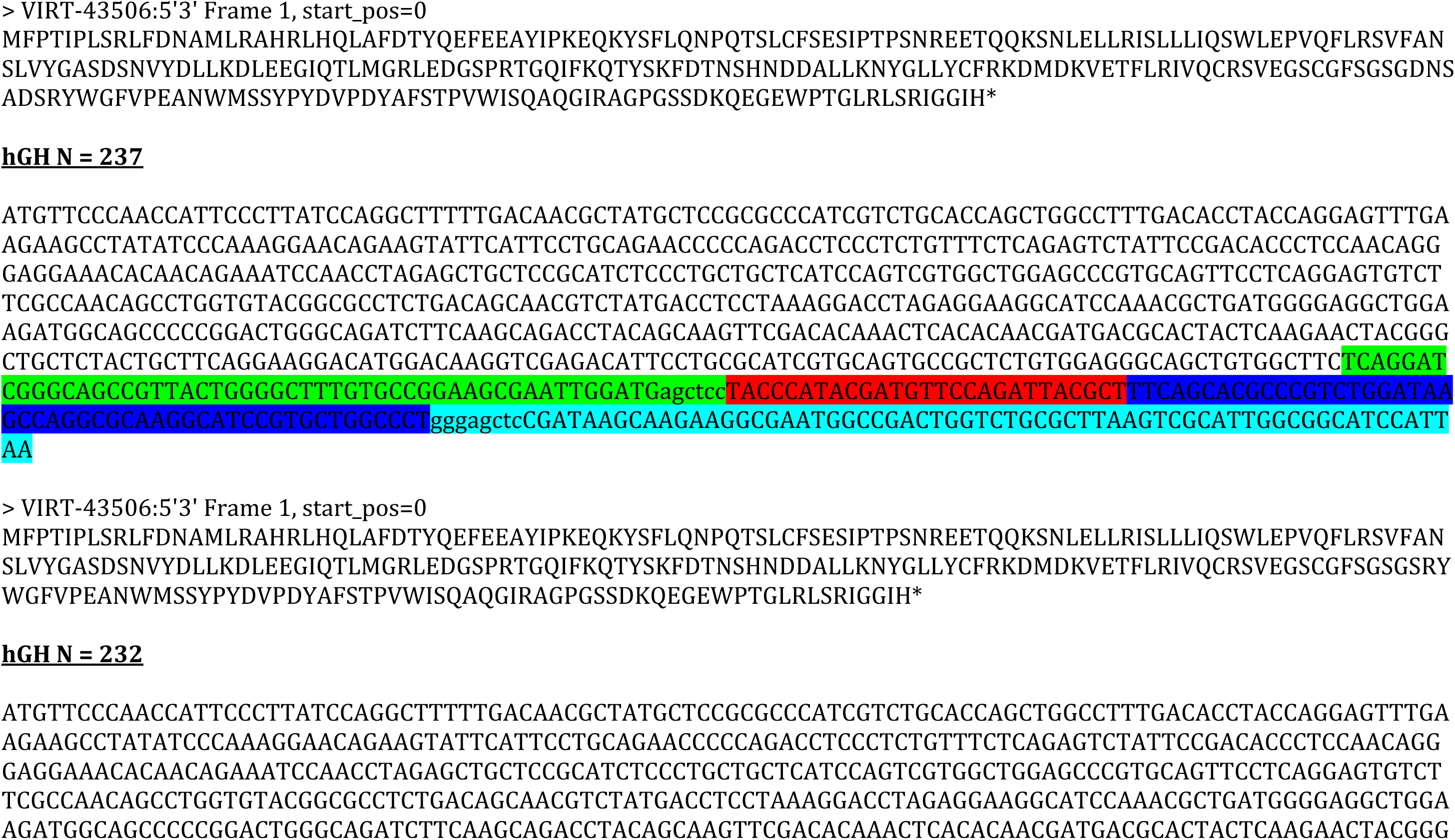
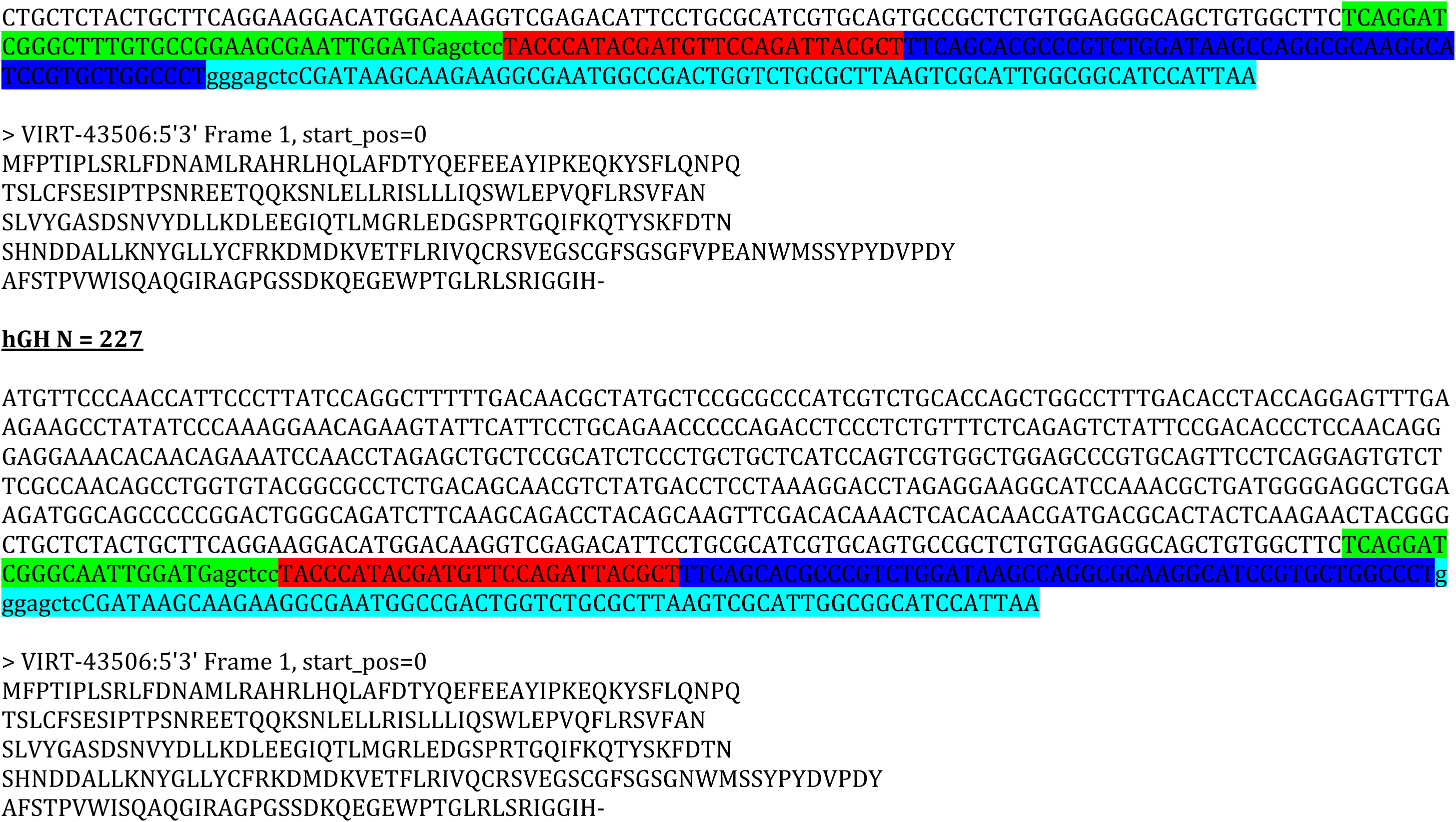
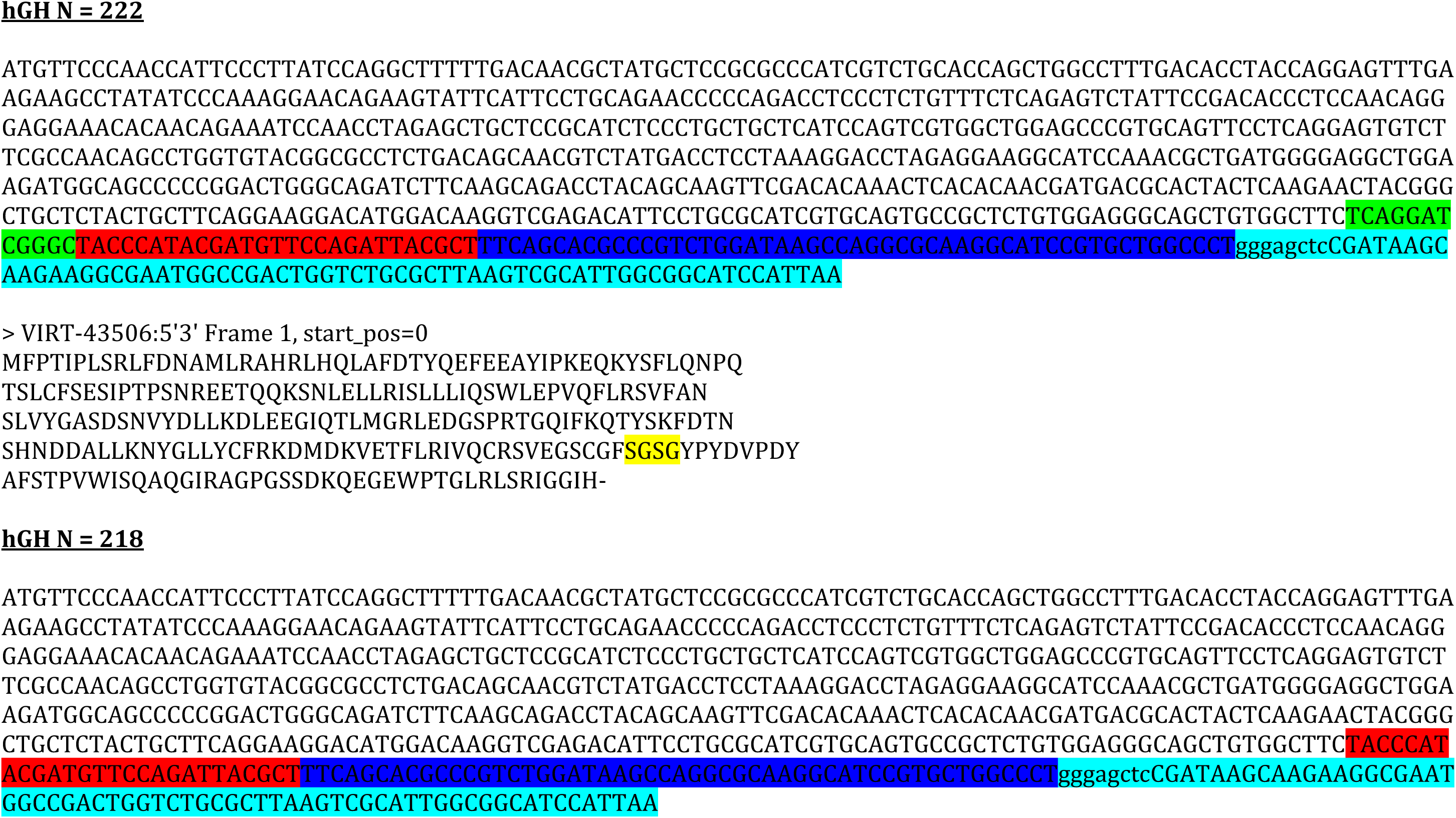
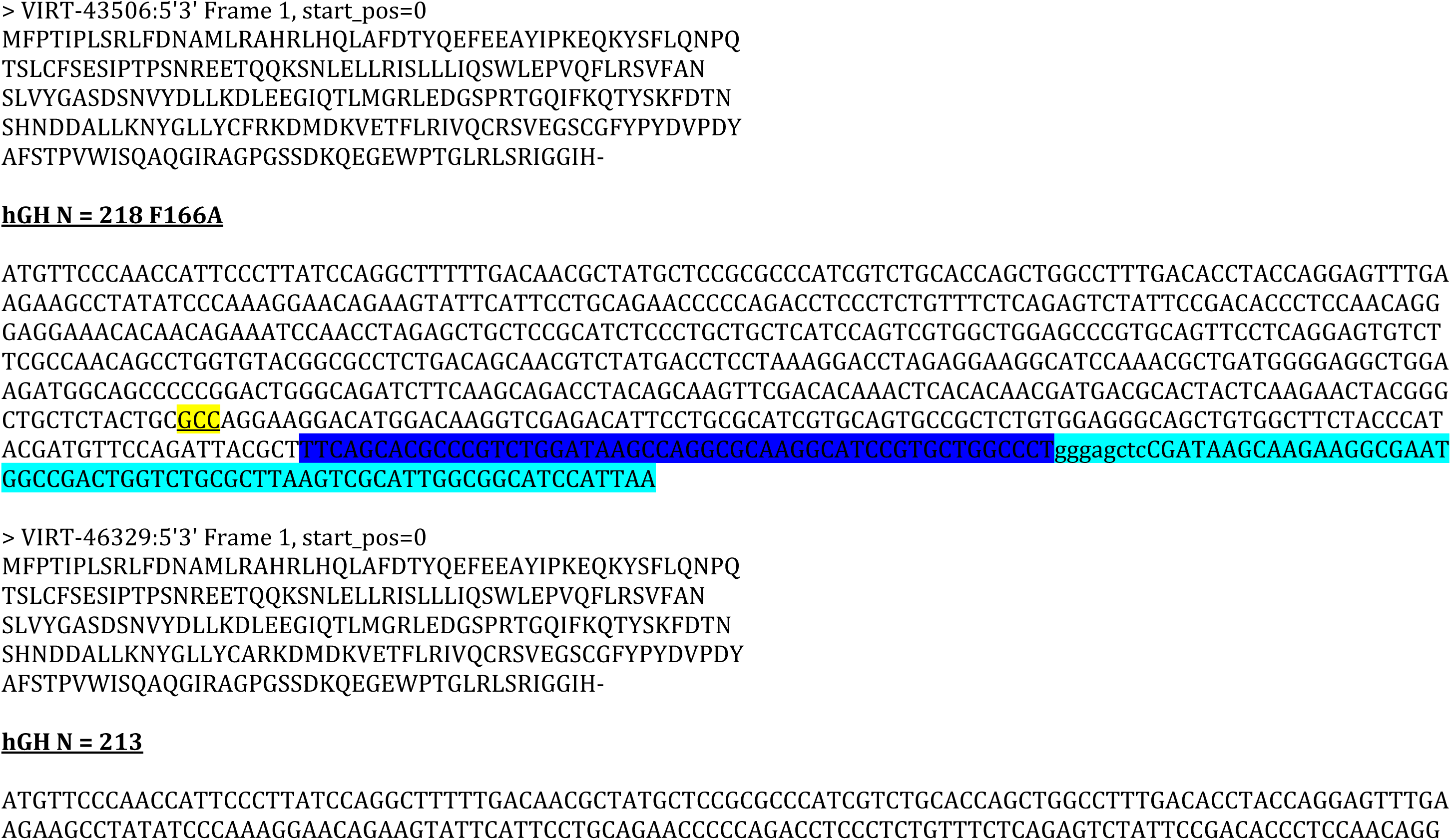
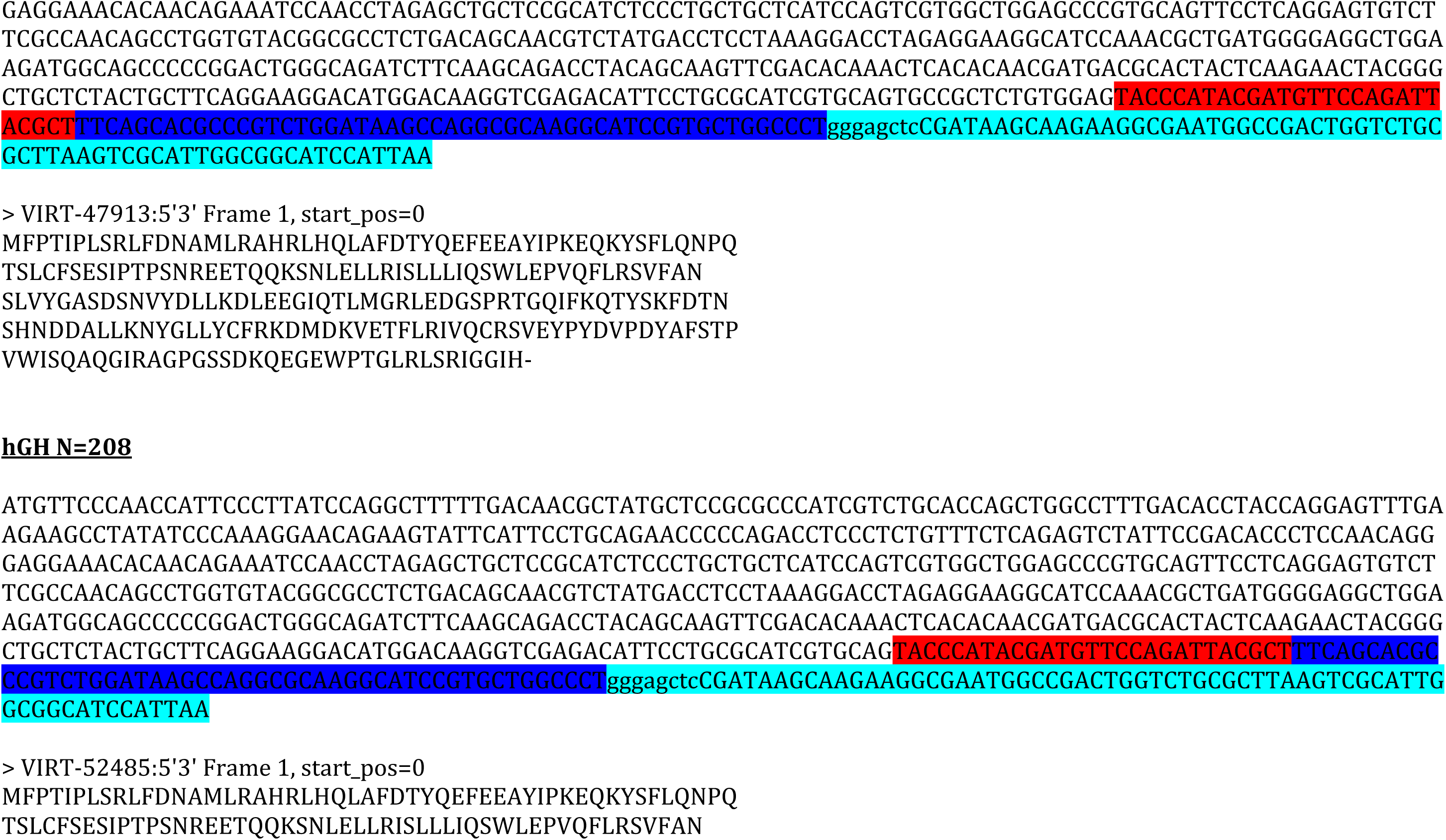
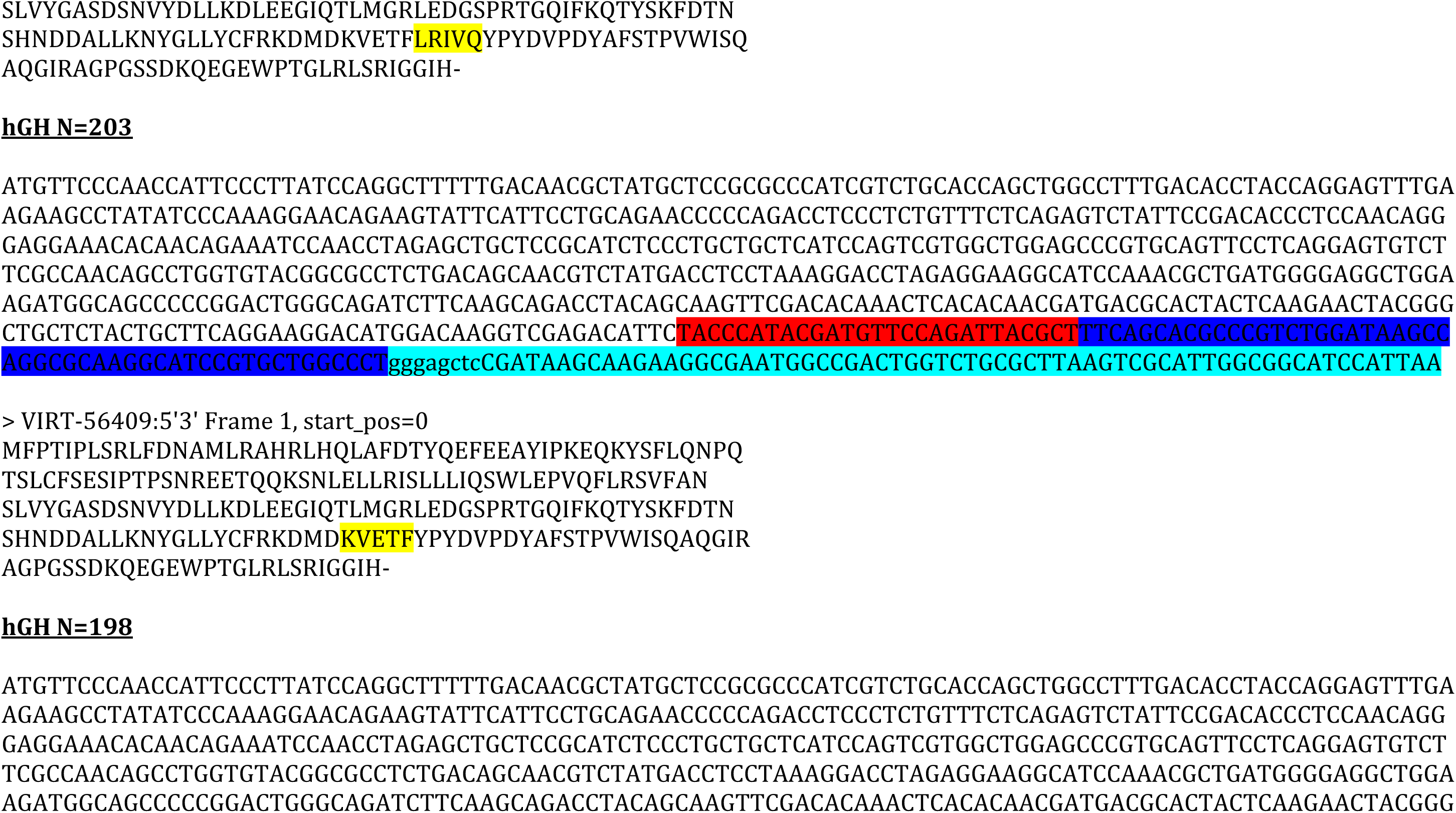
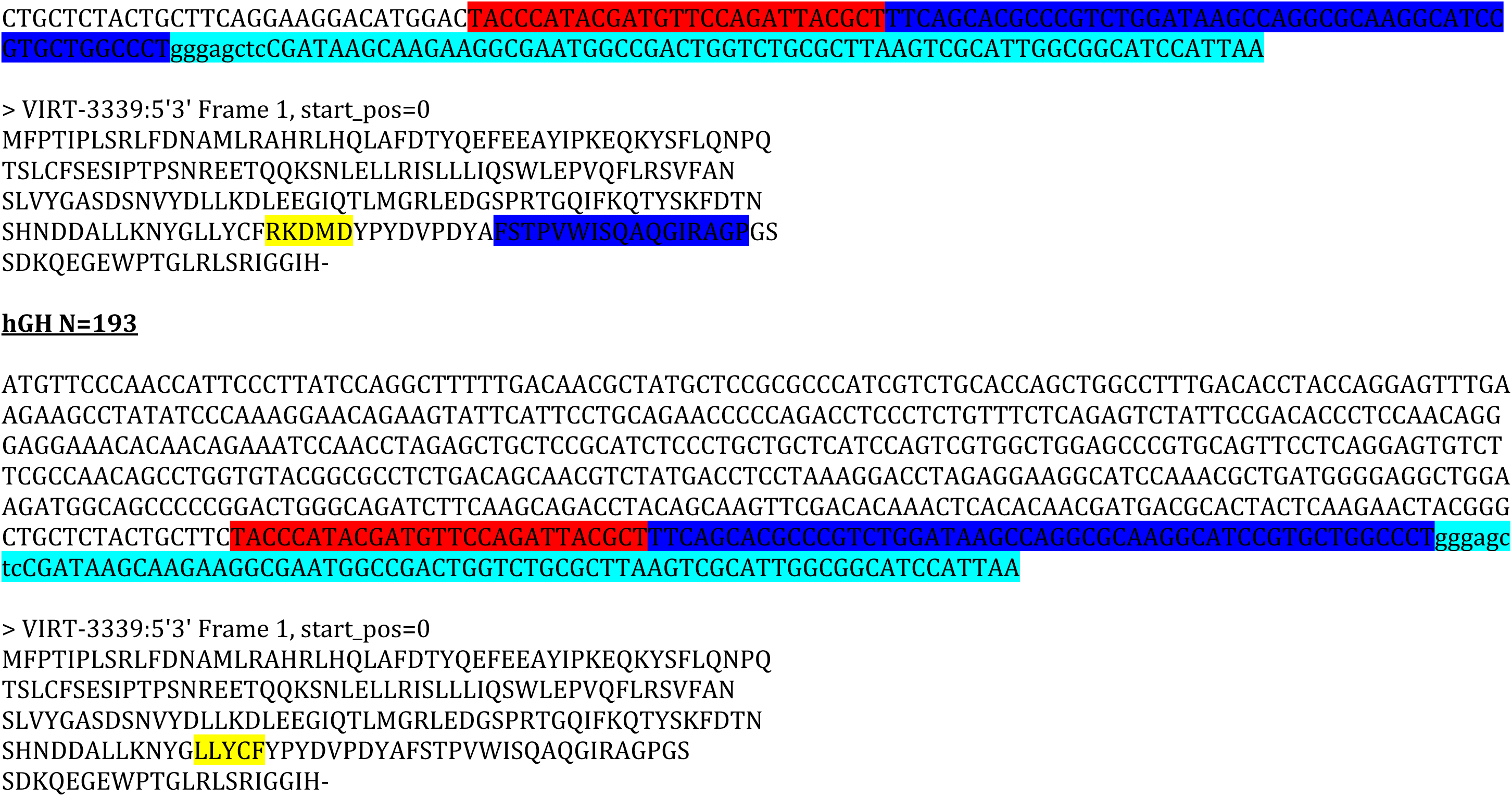
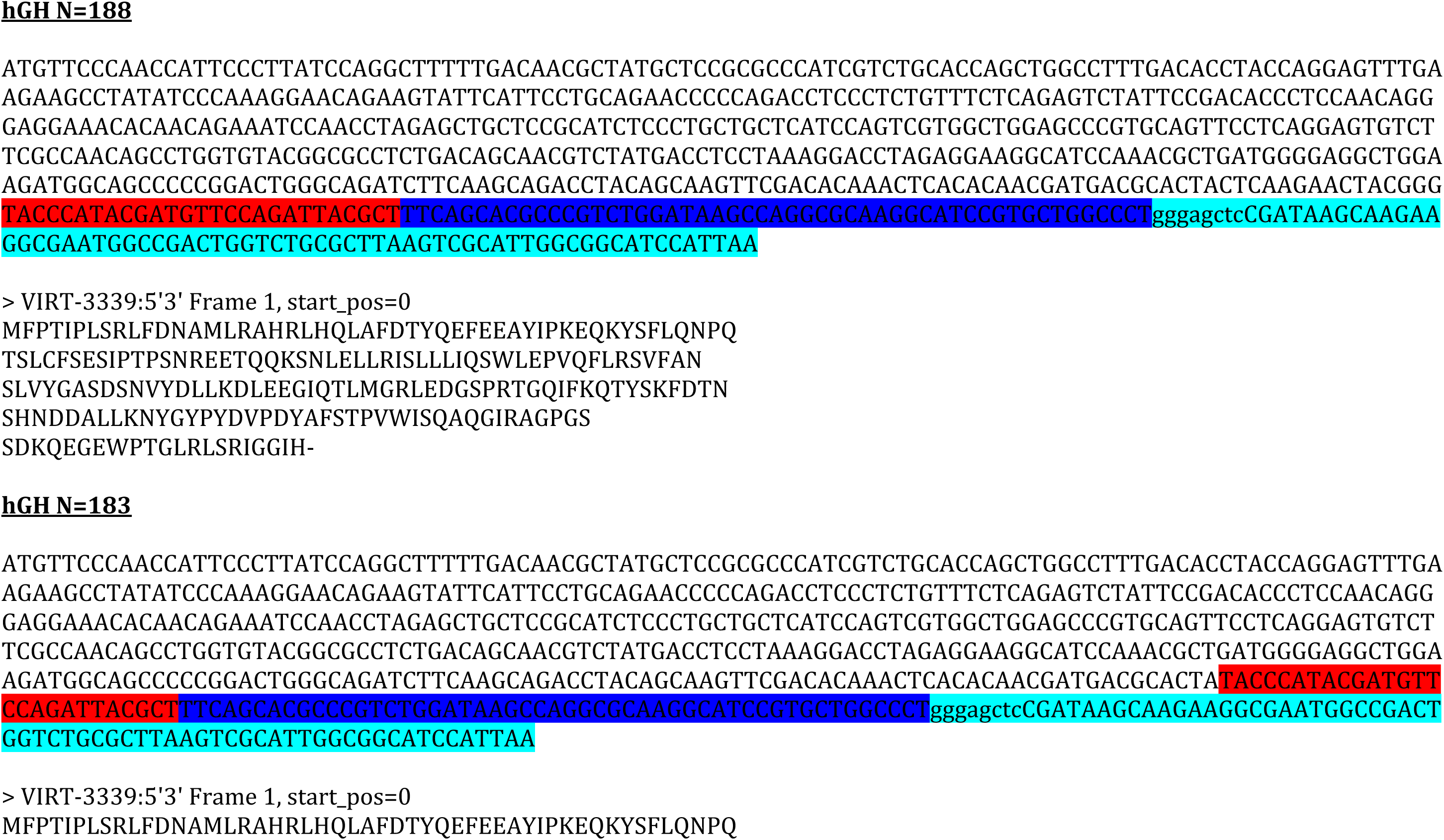
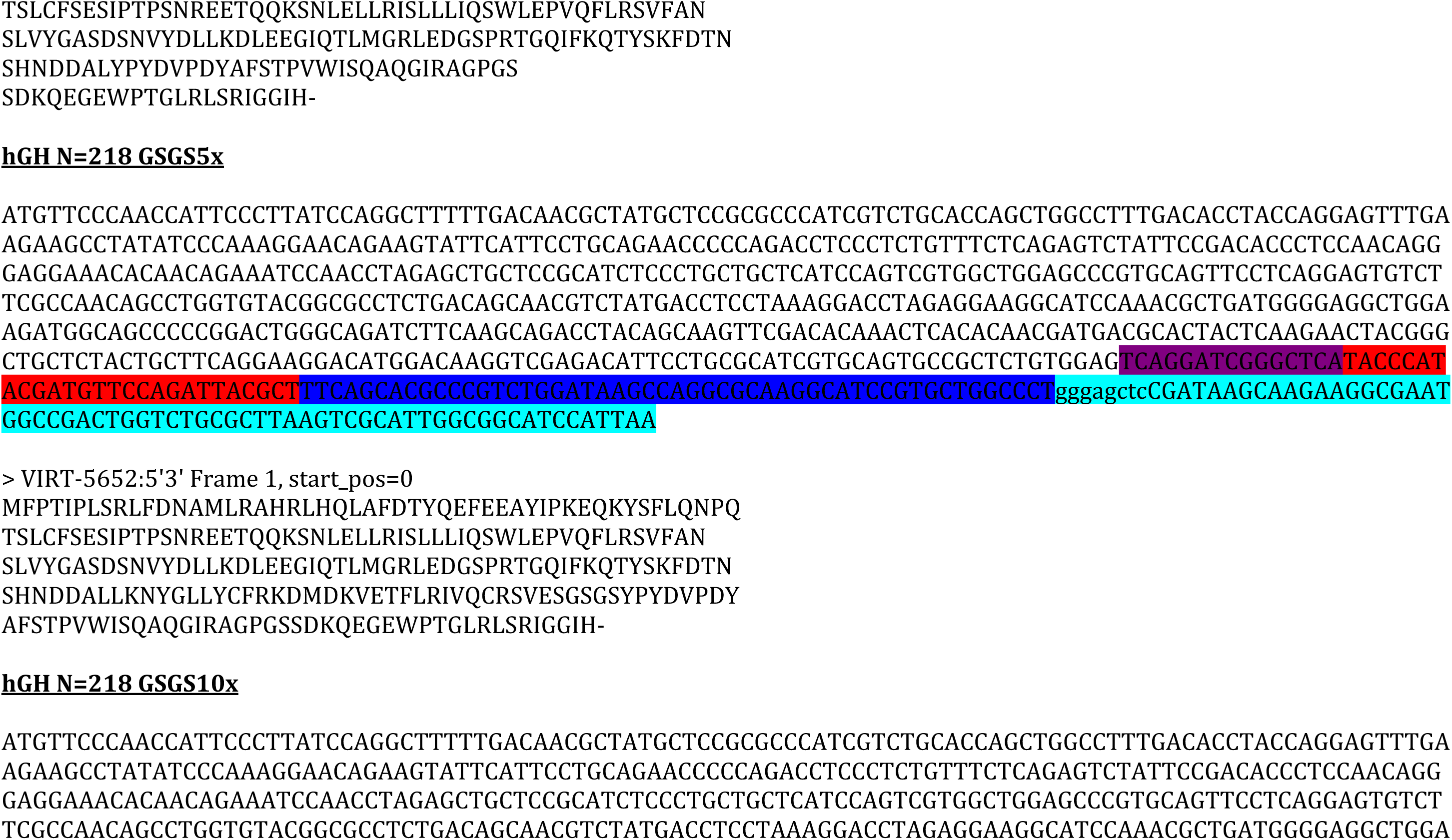
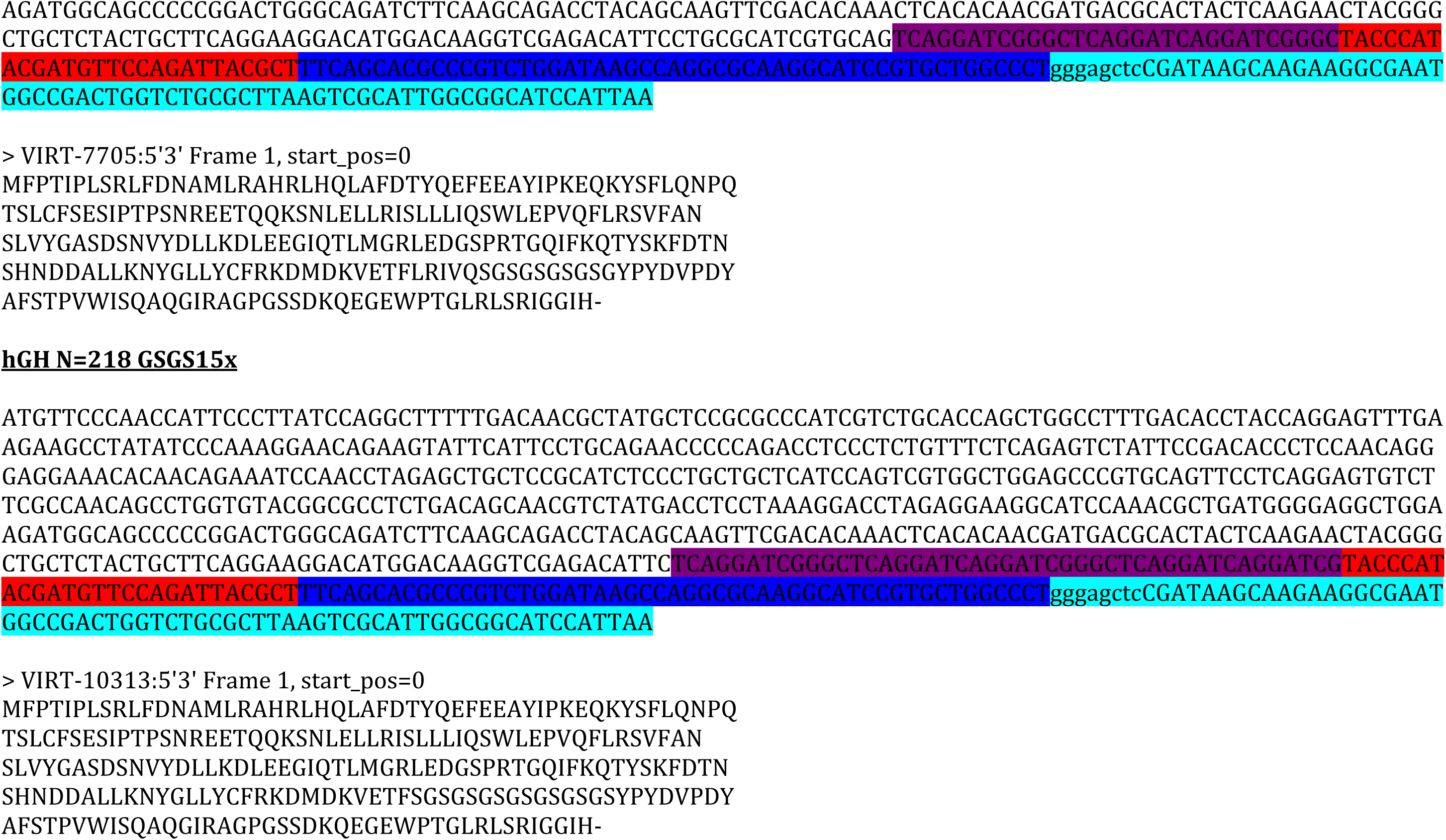
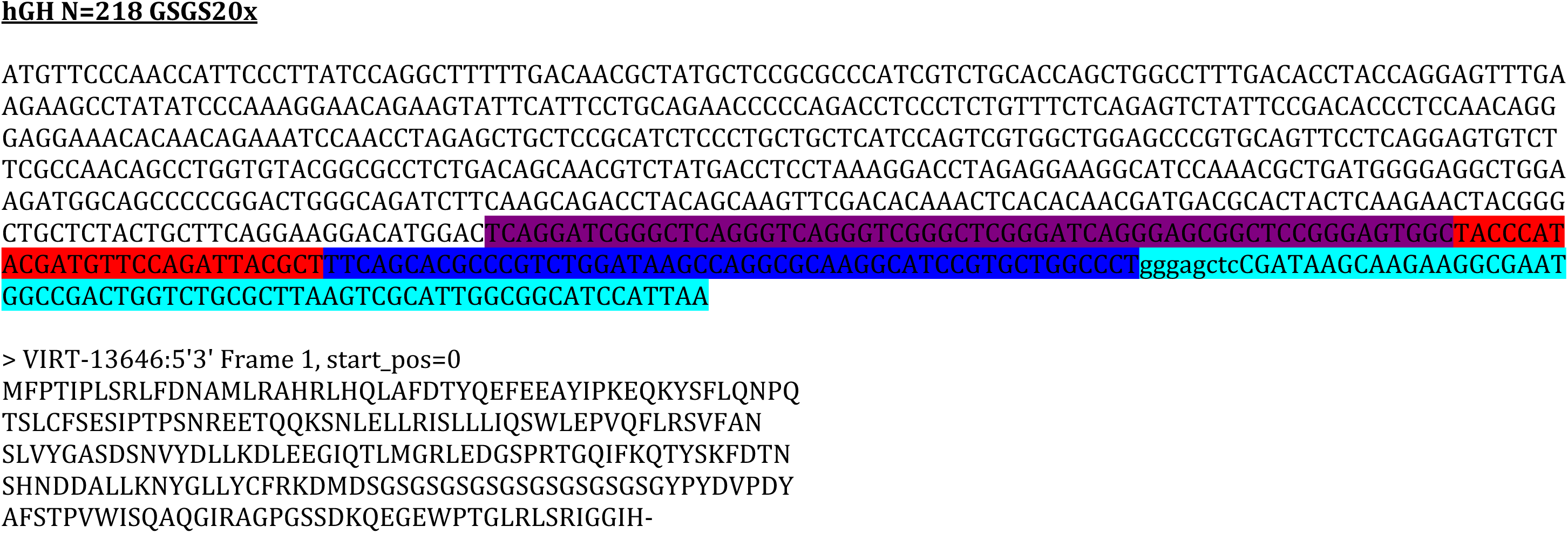
Nucleotide and amino acid sequences of all constructs

## References

[1] Ranke, M.B. and Wit, J.M. (2018). Growth hormone - past, present and future. Nat Rev Endocrinol 14, 285–300. 10.1038/nrendo.2018.22

[2] Graber, E., Reiter, E.O. and Rogol, A.D. (2021). Human Growth and Growth Hormone: From Antiquity to the Recominant Age to the Future. Front Endocrinol (Lausanne) 12, 709936. 10.3389/fendo.2021.709936

[3] Olson, K.C., Fenno, J., Lin, N., Harkins, R.N., Snider, C., Kohr, W.H. et al. (1981). Purified human growth hormone from E. coli is biologically active. Nature 293, 408–11. 10.1038/293408a0

[4] Goeddel, D.V., Heyneker, H.L., Hozumi, T., Arentzen, R., Itakura, K., Yansura, D.G. et al. (1979). Direct expression in *Escherichia coli* of a DNA sequence coding for human growth hormone. Nature 281, 544–8. 10.1038/281544a0

[5] Martial, J.A., Hallewell, R.A., Baxter, J.D. and Goodman, H.M. (1979). Human growth hormone: complementary DNA cloning and expression in bacteria. Science 205, 602–7. 10.1126/science.377496

[6] Youngman, K.M., Spencer, D.B., Brems, D.N. and DeFelippis, M.R. (1995). Kinetic analysis of the folding of human growth hormone. Influence of disulfide bonds. J Biol Chem 270, 19816–22. 10.1074/jbc.270.34.19816

[7] Nakatogawa, H. and Ito, K. (2001). Secretion monitor, SecM, undergoes self-translation arrest in the cytosol. Mol Cell 7, 185–192.

[8] Goldman, D.H., Kaiser, C.M., Milin, A., Righini, M., Tinoco, I. and Bustamante, C. (2015). Mechanical force releases nascent chain-mediated ribosome arrest *in vitro* and *in vivo*. Science 348, 457–460.

[9] Ismail, N., Hedman, R., Schiller, N. and von Heijne, G. (2012). A biphasic pulling force acts on transmembrane helices during translocon-mediated membrane integration. Nature Struct Molec Biol 19, 1018–1022.

[10] Zimmer, M.H., Niesen, M.J.M. and Miller, T.F., 3rd. (2021). Force transduction creates long-ranged coupling in ribosomes stalled by arrest peptides. Biophys J 120, 2425–2435. 10.1016/j.bpj.2021.03.041

[11] Di Palma, F., Decherchi, S., Pardo-Avila, F., Succi, S., Levitt, M., von Heijne, G. et al. (2022). Probing Interplays between Human XBP1u Translational Arrest Peptide and 80S Ribosome. J Chem Theory Comput 18, 1905–1914. 10.1021/acs.jctc.1c00796

[12] Farias-Rico, J.A., Ruud Selin, F., Myronidi, I., Frühauf, M. and von Heijne, G. (2018). Effects of protein size, thermodynamic stability, and net charge on cotranslational folding on the ribosome. Proc Natl Acad Sci U S A 115, E9280–E9287. 10.1073/pnas.1812756115

[13] Nilsson, O.B., Nickson, A.A., Hollins, J.J., Wickles, S., Steward, A., Beckmann, R. et al. (2017). Cotranslational folding of spectrin domains via partially structured states. Nat Struct Mol Biol 24, 221–225. 10.1038/nsmb.3355

[14] Farias-Rico, J.A., Goetz, S.K., Marino, J. and von Heijne, G. (2017). Mutational analysis of protein folding inside the ribosome exit tunnel. FEBS Lett 591, 155–163. 10.1002/1873-3468.12504

[15] Nilsson, O.B., Hedman, R., Marino, J., Wickles, S., Bischoff, L., Johansson, M. et al. (2015). Cotranslational protein folding inside the ribosome exit tunnel. Cell Rep 12, 1533–1540. 10.1016/j.celrep.2015.07.065

[16] Sandhu, H., Hedman, R., Cymer, F., Kudva, R., Ismail, N. and von Heijne, G. (2021). Cotranslational Translocation and Folding of a Periplasmic Protein Domain in *Escherichia coli*. J Mol Biol 10.1016/j.jmb.2021.167047. 10.1016/j.jmb.2021.167047

[17] Ismail, N., Hedman, R., Lindén, M. and von Heijne, G. (2015). Charge-driven dynamics of nascent-chain movement through the SecYEG translocon. Nat Struct Mol Biol 22, 145–149. 10.1038/nsmb.2940

[18] Nicolaus, F., Ibrahimi, F., den Besten, A. and von Heijne, G. (2022). Upstream charged and hydrophobic residues impact the timing of membrane insertion of transmembrane helices. FEBS Lett 10.1002/1873-3468.14286

[19] Nicolaus, F., Metola, A., Mermans, D., Liljenström, A., Krc, A., Abdullahi, S.M. et al. (2021). Residue-by-residue analysis of cotranslational membrane protein integration in vivo. eLife 10. 10.7554/eLife.64302

[20] Gibson, D.G., Young, L., Chuang, R.Y., Venter, J.C., Hutchison, C.A., 3rd and Smith, H.O. (2009). Enzymatic assembly of DNA molecules up to several hundred kilobases. Nat Methods 6, 343–345. 10.1038/nmeth.1318

[21] Dalbey, R.E. and Wickner, W. (1986). The role of the polar, carboxyl-terminal domain of *Escherichia coli* leader peptidase in its translocation across the plasma membrane. J Biol Chem 261, 13844–13849.

[22] Ito, K., Chiba, S. and Pogliano, K. (2010). Divergent stalling sequences sense and control cellular physiology. Biochem Biophys Res Comm 393, 1–5. 10.1016/j.bbrc.2010.01.073

[23] Butkus, M.E., Prundeanu, L.B. and Oliver, D.B. (2003). Translocon “pulling” of nascent SecM controls the duration of its translational pause and secretion-responsive secA regulation. J Bacteriol 185, 6719–6722.

[24] Cymer, F. and von Heijne, G. (2013). Cotranslational folding of membrane proteins probed by arrest-peptide-mediated force measurements. Proc Natl Acad Sci U S A 110, 14640–14645. 10.1073/pnas.1306787110

[25] Nilsson, O.B., Müller-Lucks, A., Kramer, G., Bukau, B. and von Heijne, G. (2016). Trigger factor reduces the force exerted on the nascent chain by a cotranslationally folding protein. J Mol Biol 428, 1356–1364. 10.1016/j.jmb.2016.02.014

[26] Tian, P., Steward, A., Kudva, R., Su, T., Shilling, P.J., Nickson, A.A. et al. (2018). The folding pathway of an Ig domain is conserved on and off the ribosome. Proc Natl Acad Sci USA doi.org/10.1073/pnas.1810523115

[27] Kudva, R., Tian, P., Pardo-Avila, F., Carroni, M., Best, R.B., Bernstein, H.D. et al. (2018). The shape of the bacterial ribosome exit tunnel affects cotranslational protein folding. eLife 7, e36326. 10.7554/eLife.36326

[28] Ahn, M., Wlodarski, T., Mitropoulou, A., Chan, S.H.S., Sidhu, H., Plessa, E. et al. (2022). Modulating co-translational protein folding by rational design and ribosome engineering. Nat Commun 13, 4243. 10.1038/s41467-022-31906-z

[29] Shimizu, Y., Kanamori, T. and Ueda, T. (2005). Protein synthesis by pure translation systems. Methods 36, 299–304. 10.1016/j.ymeth.2005.04.006

[30] Leininger, S.E., Rodriguez, J., Vu, Q.V., Jiang, Y., Li, M.S., Deutsch, C. et al. (2021). Ribosome Elongation Kinetics of Consecutively Charged Residues Are Coupled to Electrostatic Force. Biochemistry 60, 3223–3235. 10.1021/acs.biochem.1c00507

